# Identifying maintenance hosts and risk factors for infection with *Dichelobacter nodosus* in free-ranging wild ruminants in Switzerland: a prevalence study

**DOI:** 10.1101/691600

**Authors:** Gaia Moore-Jones, Flurin Ardüser, Salome Dürr, Stefanie Gobeli Brawand, Adrian Steiner, Patrik Zanolari, Marie-Pierre Ryser-Degiorgis

**Affiliations:** Centre for Fish and Wildlife Health, Vetsuisse-Faculty, University of Bern, Bern, Switzerland; Clinic for Ruminants, Vetsuisse-Faculty, University of Bern, Bern, Switzerland; Veterinary Public Health Institute, Vetsuisse-Faculty, University of Bern, Liebefeld, Switzerland; Institute of Veterinary Bacteriology, Vetsuisse-Faculty, University of Bern, Bern, Switzerland; Federal Food Safety and Veterinary Office (FSVO)

## Abstract

Footrot is a worldwide economically important, painful, contagious bacterial hoof disease of domestic and wild ungulates caused by *Dichelobacter nodosus* (*D. nodosus)*. Benign and virulent strains have been identified in sheep presenting mild and severe lesions, respectively. However, in Alpine ibex (*Capra ibex*) both strains have been associated with severe, fatal lesions. In Switzerland a nationwide footrot control program for sheep focusing on the virulent strain shall soon be implemented. The aim of this cross-sectional study was to estimate the nationwide prevalence of both strains of *D. nodosus* in four wild indigenous ruminant species and to identify potential susceptible maintenance hosts that could be a reinfection source for sheep. During two years (2017-2018), interdigital swabs of 1,821 wild indigenous ruminants species (Alpine ibex; Alpine chamois, *Rupicapra rupicapra*; roe deer, *Capreolus capreolus*; red deer, *Cervus elaphus*) were analysed by Real-Time PCR. Furthermore, observed interspecies interactions were documented for each sample. Overall, we report a low prevalence of *D. nodosus* in all four indigenous wild ruminants, for both the benign (1.97%, N = 36, of which 31 red deer) and virulent (0.05%, N = 1) strain of *D. nodosus.* Footrot lesions were documented in one ibex with the virulent strain, and in one ibex with the benign strain. Interactions with domestic livestock occurred mainly with cattle and sheep. In conclusion, the data suggest that domestic ungulates represent a significant source of infection for wild ungulates, while wild ruminants are likely irrelevant for the maintenance and spread of *D. nodosus*. Furthermore, we add evidence that both *D. nodosus* strains can be associated with severe disease in Alpine ibex. These data are crucial for the upcoming nationwide control program and reveal that wild ruminants should not be considered as a threat to footrot control in sheep in this context.

## Introduction

Footrot is an economically important, painful, contagious bacterial disease that affects hooves of both domestic and wild ungulates (1,2). Among domestic ruminants, mainly sheep are affected. Footrot is endemic in sheep flocks worldwide, including Switzerland (3). In free-ranging wildlife, footrot has been documented in Alpine ibex (*Capra ibex*) and mufflon (*Ovis orientalis*) (1,4). Since the treatment of wild ungulates in the field is challenging, expensive, and neither practicable nor desired by wildlife managers, severe lesions do not just affect animal welfare but typically result in death (4,5). Considering that Alpine ibex in Switzerland have just recovered from the verge of extinction in the 20^th^ century (6), outbreaks of footrot may be relevant to species conservation.

The main etiologic agent of footrot is *Dichelobacter nodosus* (*D. nodosus*) (7). Lesions begin as a mild interdigital inflammation and may progress to a separation of the horn from the underlying skin (8). Benign and virulent strains have been identified in sheep presenting mild and severe foot lesions, respectively (9). However, in Alpine ibex the benign strain was also found in association with severe lesions (4). Furthermore, both strains of *D. nodosus* have been detected in absence of lesions in sheep, cattle, goats and swine (10–13), suggesting the existence of species-specific differences in disease susceptibility and the involvement of other animal-individual or environmental risk factors in the disease course. Among others, bacterial invasion is favored by interdigital skin damage caused by trauma or environmental conditions such as humid and wet pastures (14). Unusually mild and wet weather conditions contribute to the persistence of the bacteria in the environment (1,15).

Due to an absence of long-term immunity, reinfections and clinical relapses are frequent in sheep (16). Healthy carriers are known to occur in sheep, cattle and goats (10,11,13,17) and the bacteria can survive up to two weeks in the environment (10). Virulent isolates may persist up to 10 months on bovine feet after co-grazing with sheep (11). Furthermore, a single grazing season without contact with sheep is insufficient to eliminate all pathogenic isolates from cattle feet (11). Thus, *D. nodosus* may be maintained in domestic ruminants even in the absence of clinical disease.

In the Swiss Alps, and to a lesser extent in the Jura Mountains, transhumance-grazing is a common practice and interactions between wild and domestic ruminants regularly occur on summer grazing pastures (18–20). Pastures are often shared not only among cattle, sheep, goats and South American camelids (*Lama glama* and *Vigucna pacos*) (21), but also with indigenous free-ranging wild ruminants such as ibex (*Capra ibex ibex*), chamois (*Rupicapra rupicapra*), red deer (*Cervus elaphus*) and roe deer (*Capreolus capreolus*) (20). All of them represent potential hosts of *D. nodosus* and may play a role in pathogen maintenance and spread. Based on the widespread occurrence of the disease in sheep and its sporadic character in ibex and mufflon, it has been postulated that sheep are the source of infection for wildlife, with transmission occurring on alpine pastures during the summer grazing season (1).

Currently, countrywide prevalence estimations for *D. nodosus* are available only for potential domestic hosts (13), while information on wildlife is virtually nonexistent. Importantly, further insight in the processes governing the epidemiology of *D. nodosus* infections in wild and domestic ruminants and into the infection dynamics at the wildlife-livestock interface are needed to propose appropriate disease prevention and management measures in both domestic and wild animals. For this purpose, it is crucial that data obtained on wildlife and domestic animals can be directly compared (22), which requires the use of harmonized methods, from the sampling strategy to laboratory analyses (23). Therefore, a nationwide survey on infections with *D. nodosus* in free-ranging ungulates was conducted, following the same study design and methodology as a parallel study in domestic livestock and South-American camelids in Switzerland (13).

The aim was to estimate the prevalence of *D. nodosus* infections, distinguishing between benign and virulent strains, in four species of indigenous wild ruminants during a two-year period (2017-18) in all of Switzerland. We hypothesized that wild ungulates do not maintain *D. nodosus* and that domestic ruminants act as the main infection source for wildlife. The obtained results enabled us to provide information on the likely role of the studied species in the epidemiology of footrot.

## Material & Methods

### Ethics statement

This study did not involve purposeful killing of animals. All samples originated from dead wildlife legally hunted during hunting seasons, found dead, or legally shot because of severe debilitation. According to the Swiss legislation (992.0 hunting law and 455 animal protection law, including the legislation on animal experiments; www.admin.ch), no ethical approval or permit for carcass /sample collection or animal experimentation was required.

### Study area, species of interest, study design and sampling strategy

The study area comprised the whole territory of Switzerland (41,285 km^2^), which consists of three main bioregions: the Jura Mountains (a limestone mountain chain with an elevation up to 1,679m that separates the Alps to the southwest and forms an arc to the northeast), the Midlands (characterized by a low altitude and a high human population density), and the Alps (having the highest elevation up to 4,632m above sea level, which creates a climate wall separating the south from central Europe). Species of interest were indigenous free-ranging wild ruminants including roe deer, red deer, chamois and ibex. Deer species are mostly found below the timberline - roe deer mainly in the foothill zone and red deer in the mountain zone; chamois live between the subalpine and alpine zone; and ibex above the timberline.

The study was based on a cross-sectional convenient sampling strategy with the aim of estimating a nationwide prevalence of *D. nodosus* infections on animal level in each of the four wild ruminant species. Sampling was carried out from August 2017 to December 2018, with most samples collected during the hunting seasons (i.e. August-December) of 2017 and 2018.

Sample size was calculated for each species separately, assuming simple random sampling. The free online tool by AusVet Animal Health Services (http://epitools.auvet.com) was used for the calculation, and in there, the method for the estimation of true prevalence using imperfect diagnostic test characteristics was applied. For all species the design prevalence was set at 50% because no prior information on prevalence of infection was available. The precision was set at 5% and the level of confidence at 95%. The following estimated population sizes were used for the sample size calculation (24): roe deer (113,000), red deer (28,500), chamois (91,500) and ibex (15,500). Diagnostic test characteristics (qPCR) were estimated to a mean sensitivity of 93.8% and a specificity of 98.3% (25). We aimed to sample 440-451 animals per species over two years, i.e. 1,786 individuals in total, distributed across the country and local political borders (cantons), according to species occurrence and hunting plans.

### Sample collection and animals

Sampling was mainly done by professional game wardens and hunters and in a few cases by the first and the second author. Sampling kits included one SV Lysis buffer (4 M guanindinethiocyanate, 0.01 M Tris-HL, β-mercaptoethanol) in a tube with a screw-on lid, a cotton swab, a pair of latex gloves and a data sheet to record biological data of the animal sampled (sex, age, geographical origin, body condition) and the presence of foot lesions. Documentation of foot lesions were recorded for each foot separately, including lesions in the interdigital space, the hooves and the carpal area based on the most common lesions that are used in footrot scoring systems (26). If lesions were present, the sample submitters were asked to submit the affected feet in addition to the swabs.

The study participants were also asked to report observed interspecies interactions involving both wild and domestic species within a radius of 5 km around the sampling location. This distance was roughly estimated considering the home range size of the investigated species (27,28). Four categories of contacts were used as previously described (20): physical contact; encounter of less than 50 m distance between animals; encounters of more than 50 m; and non-simultaneous occupation of the same area. These categories were not relevant for *D. nodosus* transmission but corresponded to those used in earlier studies and helped specifying the notion of interaction for the respondents. Each data sheet included two tables, one for contacts among wild ruminants and the other one for contacts between wild and domestic ungulates. For each type of contact, the frequency of observation was recorded as follows: 0) never, 1) no more than once a year, 2) more than once a year.

All sampling kits were sent to the cantonal hunting authorities, which subsequently distributed the material to game wardens or hunters who then collected the samples in the field from hunted game. Animals that were sent as routine diagnostic cases (without footrot lesions) to the Centre for Fish and Wildlife Health (FIWI, University of Bern) for post-mortem examination were also sampled for this study (N = 54).

A four-feet sample was taken from each animal, i.e. the interdigital space of each of the four feet of each animal was sampled with the same cotton swab (2 mm 15 cm, Heinz Herenz, Medizinalbedarf GmbH, Hamburg, Germany). Each clean quarter of the swab was used for each foot as previously described (17), i.e. there was one swab per animal for laboratory analysis.

Immediately after swabbing the feet, each swab was immersed for at least 1 min in a tube containing SV lysis buffer with a hermetic lid. This procedure allowed transportation and storage without cooling until analysis (9).

In total 1,821 samples were taken, of which 91.4% (N = 1,664) by game wardens/hunters; 4.5% (N = 83) by field biologists; 3.3% (N = 60) by veterinarians (FIWI) and for 0.7% (N = 14) of the samples the profession of the sampler was not given. The feet of five ibex, one red deer, three chamois and two roe deer with presumptive footrot lesions were sent to the FIWI for macroscopic examination. All 26 cantons contributed to the achieved sample number. Thirty-seven point seven percent of the samples (N = 686) were received in 2017 and 62.3% (N = 1135) in 2018.

There were 961 males, 807 females and 53 animals without sex information. We used three age categories for the age estimation: juvenile (< 1 year, N=167), yearling (1-<2 year; N=245) and adult (≥ 2 years, N=1,335). Age estimation was based on acquired knowledge of the sample submitters during their formal training (hunters and professional game wardens). For roe deer and red deer this was based on the tooth replacement and wear, while for ibex and chamois it was based on the growth rings of the horns (29). No age information was provided for 74 animals.

### Laboratory analysis

All collected samples were tested for the presence of benign or/and virulent strains of *D. nodosus* by PCR (9). Deoxyribonucleic acid (DNA) was extracted from the buffer solution by an automated method using a KingFisher™ Duo Prime purification automat. A competitive real-time PCR was performed in all samples, which allows the simultaneous detection of the virulent and benign strains of *D. nodosus* by distinguishing the extracellular protease AprV2 found in the more virulent strains from the subtly different protease AprB2 that is found in strains associated with benign disease signs in sheep (9). Amplification was done in a 7500 Real-Time PCR system with cycle conditions of 2 min at 50° C, 10 min at 95° C followed by 40 cycles of 15s at 95°C and 1 min at 60° C. Samples that showed no fluorescent signal were considered negative; samples showing a positive fluorescent signal for either strain were classified as positive. Results were then analyzed using the Sequence Detector 7500.

### Data management and statistical analysis

Data handling, validating, cleaning and coding were done in MS Excel spreadsheets followed by transfer to R software version 3.5.1. (https://cran.r-project.org) for all statistical analyses. Apparent and true prevalence were calculated using the “epi.prev” function of the package “epiR” considering imperfect test characteristics, with a test sensitivity of 93.8% and test specificity of 98.3% (25) and a confidence interval of 95%. The function calculates the true prevalence (TP) from the apparent prevalence (AP) using the Rogan-Gladen estimator, based in the formula *TP* = (*AP* + *specificity* − 1)/(*sensitivity* + *specificity* − 1) (30).

A risk factor analysis for presence of *D. nodosus* on wild animal feet was performed with the PCR status of the animals (positive or negative) as dependent variable and potential risk factors as independent variables. Because only one sample was positive for the virulent strain, the risk factor analysis was performed for the benign strain only. Logistic regression models were used first in a univariable analysis followed by a multivariable analysis. The two variables interactions with either roe deer or with chamois, were tested using Fisher’s Exact Test because zero interactions were observed for PCR-positive animals. Variables with p-values < 0.2 were further considered in the multivariable analysis, where a manual backward elimination procedure was performed with a cut-off level at p-value <0.05. To be able to compare the prevalence for the benign strain of *D. nodosus* in sheep and cattle (domestic ungulate study; Ardüser et. al 2019) with the prevalence of infection for the benign strain in all wild ungulates (all species pooled; this study), we used a logistic regression model in a multivariable analysis with a cut-off level at a p-value <0.05.

All maps were generated with QGIS 2.18.16 Las Palmas (Free Software Foundation Inc., Boston, USA).

## Results

### Detection of *Dichelobacter nodosus*

Out of 1,821 sampled animals (Figure 1), 37 were found positive for *D. nodosus* (Figure 2). The virulent strain was only found in one animal, an adult male ibex with severe disease signs. The benign strain was found in 36 animals, including 31 red deer (20 males and 11 females; 8 juveniles, 5 yearlings, 17 adults, and 1 animal of unknown age), one roe deer (yearling female), one Alpine chamois (adult male) and three Alpine ibex (all adult males). The estimated apparent prevalence for the benign and virulent strains by species is given in Table 1a. Due to the low prevalence of *D. nodosus* per species, the true prevalence (TP) could only be calculated for red deer (TP= 6.08% CI=3.5 – 9.3).

**Table 1a.** Prevalence of *D. nodosus* in wild ungulates.

| Species | Tested samples | Positive benign | Benign %<br>AP | Positive virulent | Virulent %<br>AP |
| --- | --- | --- | --- | --- | --- |
| Alpine ibex | 589 | 3 | 0.50%<br>(0.13 - 1.46) | 1 | 0.16 %<br>(0.00 - 0.94) |
| Alpine chamois | 410 | 1 | 0.24%<br>(0.00 – 1.35) | 0 | 0.00%<br>(0.00 – 0.87) |
| Red deer | 408 | 31 | 7.59%<br>(5.29 – 10.58) | 0 | 0.00%<br>(0.00– 0.88) |
| Roe deer | 409 | 1 | 0.24%<br>(0.00 – 1.35) | 0 | 0.00%<br>(0.00 - 0.87) |
| TOTAL | 1821* | 36 | 1.97%<br>(1.39 – 2.72) | 1 | 0.05%<br>(0.00 – 0.30) |

Number of samples from wild ungulates, PCR results and estimated apparent prevalence (AP) with corresponding 95% confidence intervals indicated in parentheses.
\*For five samples, the species was not indicated by the submitter.

**Table 1b.** Prevalence of *D. nodosus* in domestic ungulates for comparison.

| Species | Tested<br>samples | Positive<br>benign | Benign %<br>AP | Positive<br>virulent | Virulent %<br>AP |
| --- | --- | --- | --- | --- | --- |
| Sheep | 690 | 58 | 7.80%<br>(3.40 - 12.10) | 94 | 17.60 %<br>(10.70 - 24.40) |
| Cattle | 849 | 694 | 83.30%<br>(79.10 – 87.50) | 0 | 0.00%<br>(0.00 – 0.40) |
| Goats | 790 | 23 | 2.30%<br>(0.00 – 5.10) | 0 | 0.00%<br>(0.00– 0.50) |
| SAC | 591 | 13 | 0.90%<br>(0.30 – 1.50) | 3 | 0.20%<br>(0.00 - 0.40) |
| TOTAL | 2920 | 788 | 26.98%<br>(23.38-28.62) | 97 | 3.21%<br>(2.70-4.02) |

Ardüser et. al 2019

**Fig 1.**
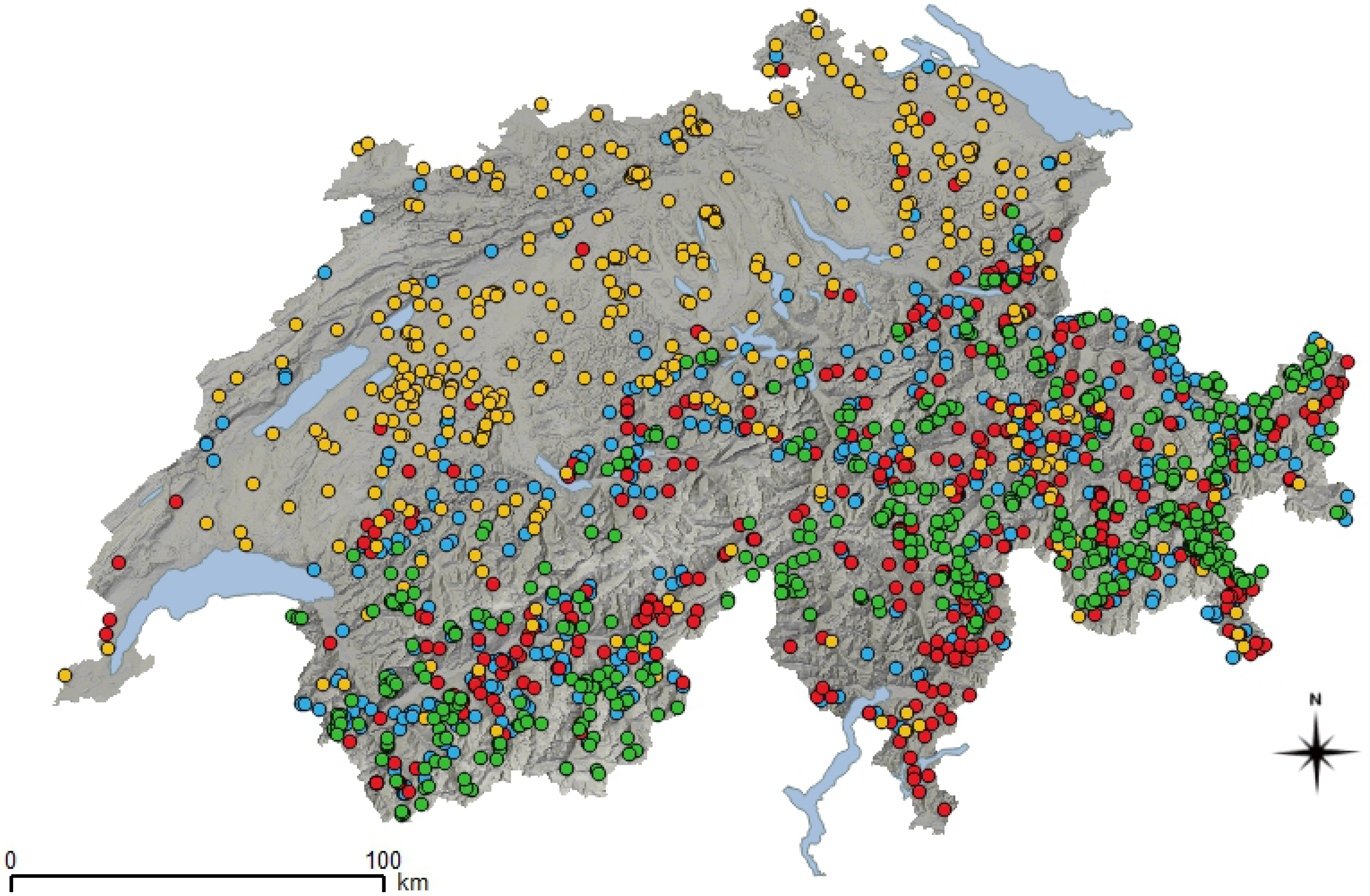
Map of Switzerland showing the distribution of the sampled wild animals. Shades of grey illustrate the relief. Lakes are in blue. Colored dots correspond to animals of different species: Yellow: roe deer; Red: red deer; Blue: chamois; and Green: ibex.

**Fig 2.**
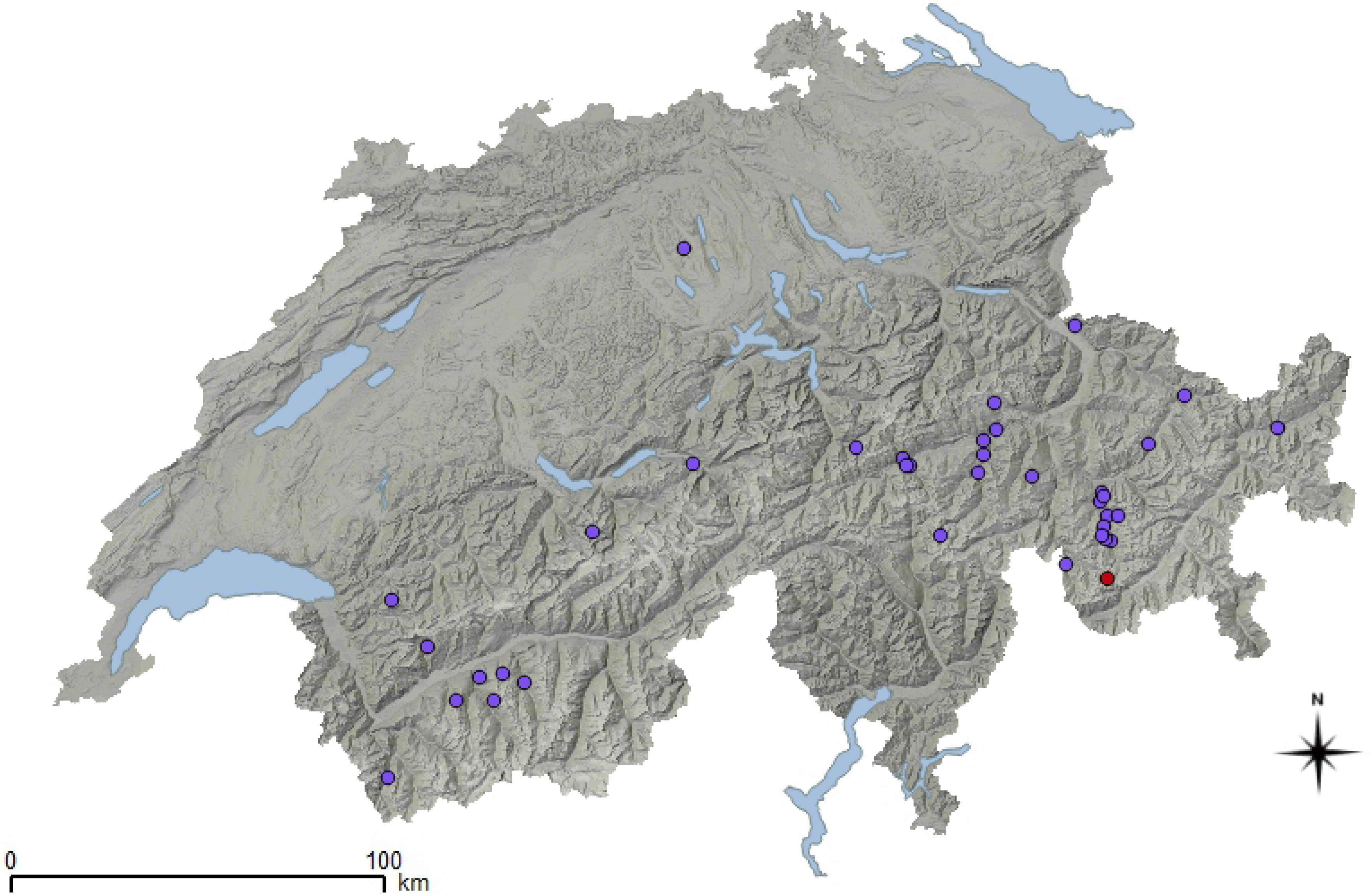
Map of Switzerland showing the geographical origin of PCR positive animals. Shades of grey illustrate the relief. Lakes are in blue. Violet dots correspond to animals positive for the benign strain and red dots to animals positive for the virulent strain.

All positive samples originated from animals from six out of 26 Swiss cantons (S1 Table) all of them located in subalpine to alpine zones (> 1000 m above sea level) except for five red deer and one roe deer from the mountain and foothill zones. The majority (N=27, 68%) of the positive animals originated from the canton of Grisons (23 red deer with the benign strain; and two ibex: one positive for the benign and the other one for the virulent strain). Positive samples were found in both sampling years (14 in 2017; 23 in 2018).

### Foot lesions

Four of the 31 (12.9%) red deer positive for the benign strain of *D. nodosus* were reported by the sample submitters as having foot lesions. Two of them presented with lesions that consisted in overgrown hooves, one of which concerned the lateral digits only, without noticeable interdigital inflammation (i.e. no signs suggestive of footrot). In the other two red deer, an interdigital grey malodorous exudate and rotting of the sole of one of the forelimbs was reported. However, these observations could not be confirmed as neither photographs nor any of the affected feet were submitted with the samples. One ibex positive for the benign strain presented severe footrot lesions in the hind feet (Figure 3). All other animals positive for the benign strain did not show any sign of disease (including two ibex).

**Fig 3.**
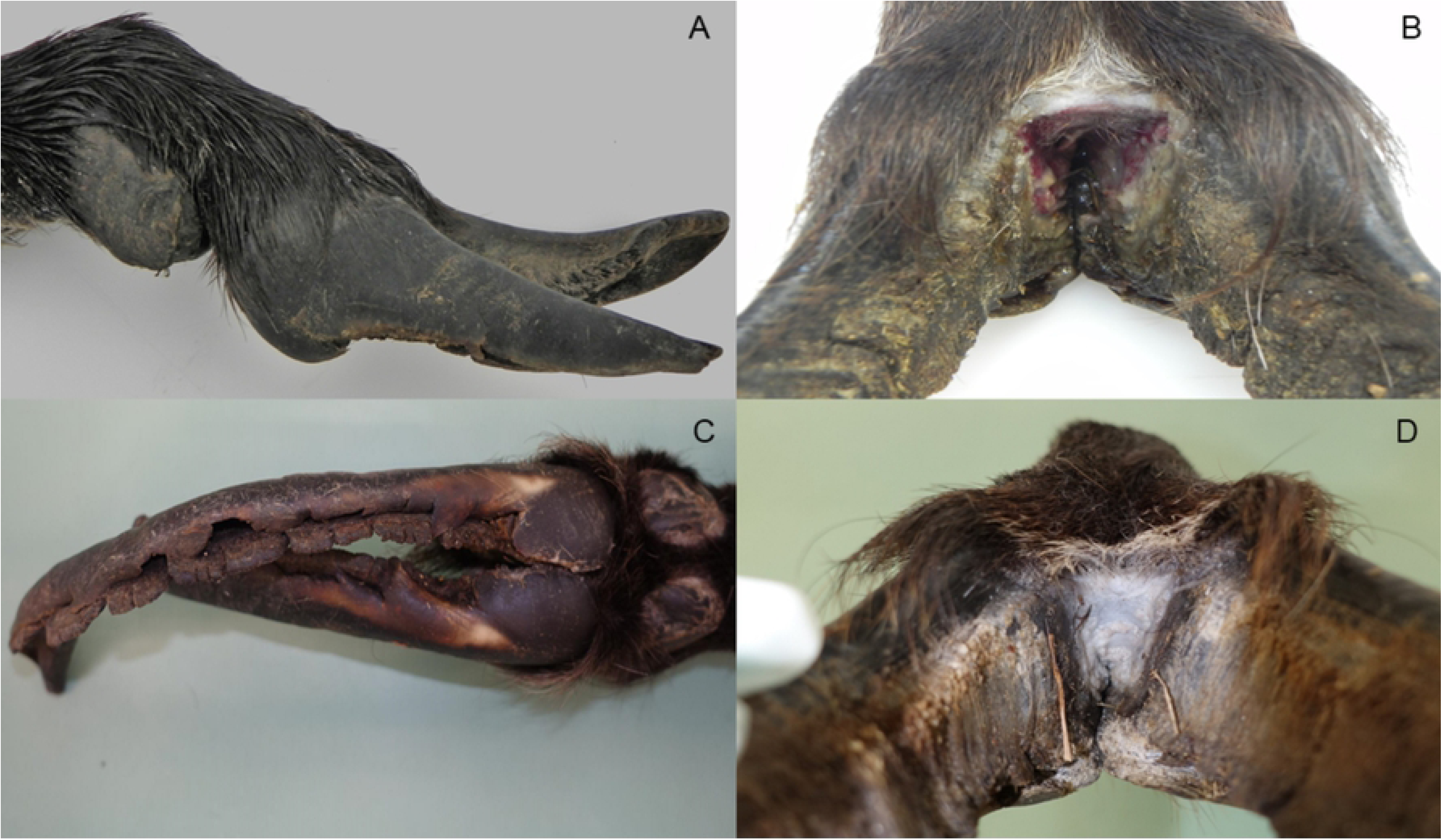
*D. nodosus* positive feet of two ibex. Front feet of an ibex positive for the virulent strain of *D. nodosus*: (A) severely overgrown hooves and (B) ulceration and greyish discoloration of the interdigital space. Hind feet of an ibex positive for the benign strain of *D. nodosus*: (C) greyish discoloration of the interdigital space and (D) severely overgrown and fissurated hooves.

The ibex with the virulent strain of *D. nodosus* showed severe footrot lesions in the front feet (Figure 3), including a severe ulcerative interdigital pododermatitis with greyish discoloration associated with an unpleasant odor, and severely multifocally fissured and overgrown hooves. The severity of the lesions were similar in the two ibex infected with the benign and the virulent strain, respectively.

### Interspecies interactions

In total, 68% (N = 1236) of the sample submitters answered the questions listed on the data sheet regarding interspecies interactions in their sampling area, either partially (58%) or fully (10%). Data on reported proximity among species are summarized in Figure 4.

**Fig 4.**
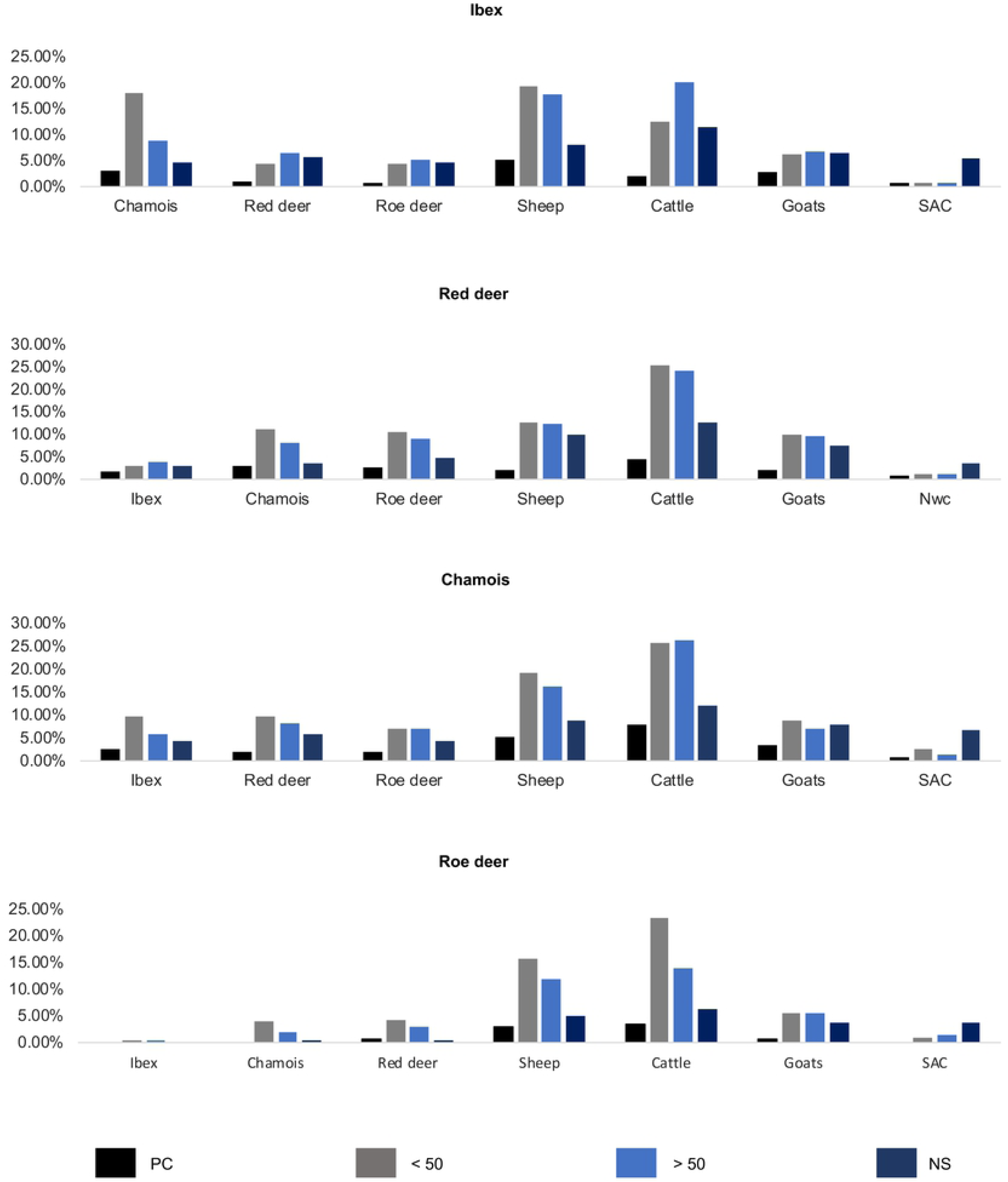
Relative frequency of observations of different types of interspecies contacts between wild and domestic ruminants in Switzerland. Proximity between species: 100% relates to all samples collected (information from respondents who did not report observations and lacking information due to non-reporting are not included in the graph). Colored bars indicate the following types of interactions: PC: physical contact); <50 m: encounter of <50 m; >50 m: encounter of >50 m; NS: non-simultaneous occupation of the same pasture.

Encounters with both wildlife and domestic species were reported for all four investigated species in all possible categories. Encounters of <50m, >50m and the non-simultaneous occupation of the same pasture were reported more frequently with cattle and sheep than with goats and South American camelids.

For ibex and chamois more specifically, about a third of the respondents reported encounters of <50m and >50m with sheep as well as with cattle in ibex. Regarding chamois only, half of the respondents reported encounters of <50m and >50m with cattle. As for the non-simultaneous occupation of pastures, 11% of the respondents mentioned the use of pastures in wildlife habitat by cattle and 8% by sheep. Physical contact between ibex and domestic livestock was not frequent and reported mainly with small domestic ruminants (8% of the respondents). For chamois, a similar percentage of physical contact was reported with small domestic ruminants (9%) and cattle (8%). As concerns red deer, encounters of <50m (25%) and >50m (24%) were observed with cattle, and 4% of the respondents even reported physical contact between the two species. Regarding roe deer, patterns of encounters of <50m (23%) and >50m (14%) with cattle were similar to the observed interactions that were reported for red deer. The non-simultaneous occupation of the same pasture by deer and domestic ungulates was reported more frequently with cattle (red deer: 13%; roe deer: 6%) than with the other three domestic species.

Considering wild species only, interspecific interactions were reported mainly within the same taxonomic family, i.e. among caprids (ibex and chamois) and among cervids (roe deer and red deer). Interactions between red deer and chamois were also frequently reported and occasionally included physical contacts (5% of the respondents). By contrast, only a few interactions between ibex and cervids were reported, even though rare physical contacts between ibex and red deer (2%) were documented.

Focusing on positive samples, 83% of the submitters of the 28 red deer and the single roe deer samples positive for the benign strain of *D. nodosus* reported having observed direct or indirect contacts with deer and cattle in the sampling area. Regarding the single positive chamois, contacts among all wild and domestic species except domestic goats were reported to have been observed in the sampling area. As concerns the four positive ibex, interspecies interactions were recorded only in the sampling area of two of them (both positive for the benign strain but without lesions), with all four domestic species and with sheep only, respectively. As for the two ibex with severe footrot lesions, no information regarding interspecies interactions was provided.

### Analysis of risk factors for infection with *D. nodosus*

For the benign strain, univariable analysis indicated that the variables sex (male *vs* female), age (adult *vs* yearling and juvenile), species (ibex *vs* red deer, chamois and roe deer), the presence of foot lesions (presence *vs* absence) and interspecies interactions in the sampling area (report *vs* no report) were associated for some levels with an infection of benign *D. nodosus* considering the threshold of p = 0.2 (Table 2).

**Table 2.** Univariable risk factor analysis for presence of benign *Dichelobacter nodosus* on wild ruminant feet.

| Variables |  | Tested/total (%) |  | p-value | OR95% |
| --- | --- | --- | --- | --- | --- |
|  |  | positive for<br>benign<br>*D. nodosus* | negative for<br>benign<br>*D. nodosus* |  |  |
| Sex | Female | 12/807<br>(1.5%) | 795/807<br>(98.5%) | - | Baseline |
|  | Male | 24/961<br>(2.5%) | 937/961<br>(97.5%) | 0.138 | 1.69 (0.85 - 3.53) |
| Age | Juvenile | 8/167<br>(4.8%) | 159/167<br>(95.2%) | - | Baseline |
|  | Yearling | 6/245<br>(2.4%) | 239/245<br>(97.6%) | 0.205 | 0.49 (0.16 - 1.46) |
|  | Adult | 21/1335<br>(1.6%) | 1314/1335<br>(98.4%) | 0.006 | 0.31 (0.14 – 0.77) |
| Species | Ibex | 3/589<br>(0.5%) | 586/589<br>(99.5%) | - | Baseline |
|  | Red deer | 31/408<br>(7.6%) | 377/408<br>(92.4%) | < 0.001 | 16.06 (5.68 - 67.24) |
|  | Chamois | 1/410<br>(0.2%) | 409/410<br>(99.8%) | 0.523 | 0.47 (0.03 - 3.74) |
|  | Roe deer | 1/409<br>(0.2%) | 408/409<br>(99.8%) | 0.524 | 0.47 (0.02 - 3.75) |
| Foot lesions | Without foot lesion | 29/1449<br>(2.0%) | 1420/1449<br>(98.0%) | - | Baseline |
|  | With foot lesion | 5/76<br>(6.5%) | 71/76<br>(93.5%) | 0.013 | 3.44 (1.14 - 8.45) |
| *Contacts | No report with cattle | 5/254<br>(1.6%) | 249/254<br>(98.4%) | - | Baseline |
|  | With cattle | 26/751<br>(3.5%) | 725/751<br>(96.5%) | 0.240 | 1.78 (0.73 - 5.32) |
|  | No report<br>with goats | 7/387<br>(1.6%) | 380/387<br>(98.4%) | - | Baseline |
|  | With<br>goats | 10/322<br>(3.1%) | 312/322<br>(96.9%) | 0.267 | 1.73 (0.66 - 4.84) |
|  | No report<br>with ibex | 1/107<br>(0.9%) | 106/107<br>(99.1%) | - | Baseline |
|  | With ibex | 7/99<br>(7.0%) | 92/99<br>(93.0%) | 0.052 | 4.01 (0.70 - 75.46) |
|  | No report<br>with<br>chamois | 0/68<br>(0.00%) | 68/68<br>(100%) | 0.123 | Inf. |
|  | With<br>chamois | 10/231<br>(4.3%) | 221/231<br>(95.7%) |  |  |
|  | No report<br>with roe<br>deer | 0/68<br>(0.00%) | 68/68<br>(100%) | 0.066 | Inf. |
|  | With roe<br>deer | 10/189<br>(5.3%) | 179/189<br>(94.7%) |  |  |

†The variable “contacts” refers to reported interspecies interactions with wild and domestic ungulates (all *D. nodosus* positive wild animal species pooled) within a radius of 5km around the sampling location.

The factor “canton” was also investigated but showed a weak association with an infection of *D. nodosus* (p = 0.68). All factors with a p value < 0.2, except “interspecies contacts” due to the large number of missing values, were additionally tested in a multivariable analysis, which revealed that the species “red deer” and the presence of foot lesions remained significant risk factors for infection (Table 3). The model showed that red deer were 15.97 times more likely to be infected with the benign strain of *D. nodosus* than ibex (CI: 5.49 - 68.17, estimate: 2.7710, standard error: 0.6209, z-value: 4.463, Pr (|z|) = < 0.001***) and that animals with foot lesions were 5.39 more likely to be infected with the benign strain of *D. nodosus* than animals without lesions (CI: 1.64 - 15.37 estimate: 1.6848, standard error: 0.5584, z-value: 3.017, Pr (|z|) = < 0.00255**).

**Table 3.**
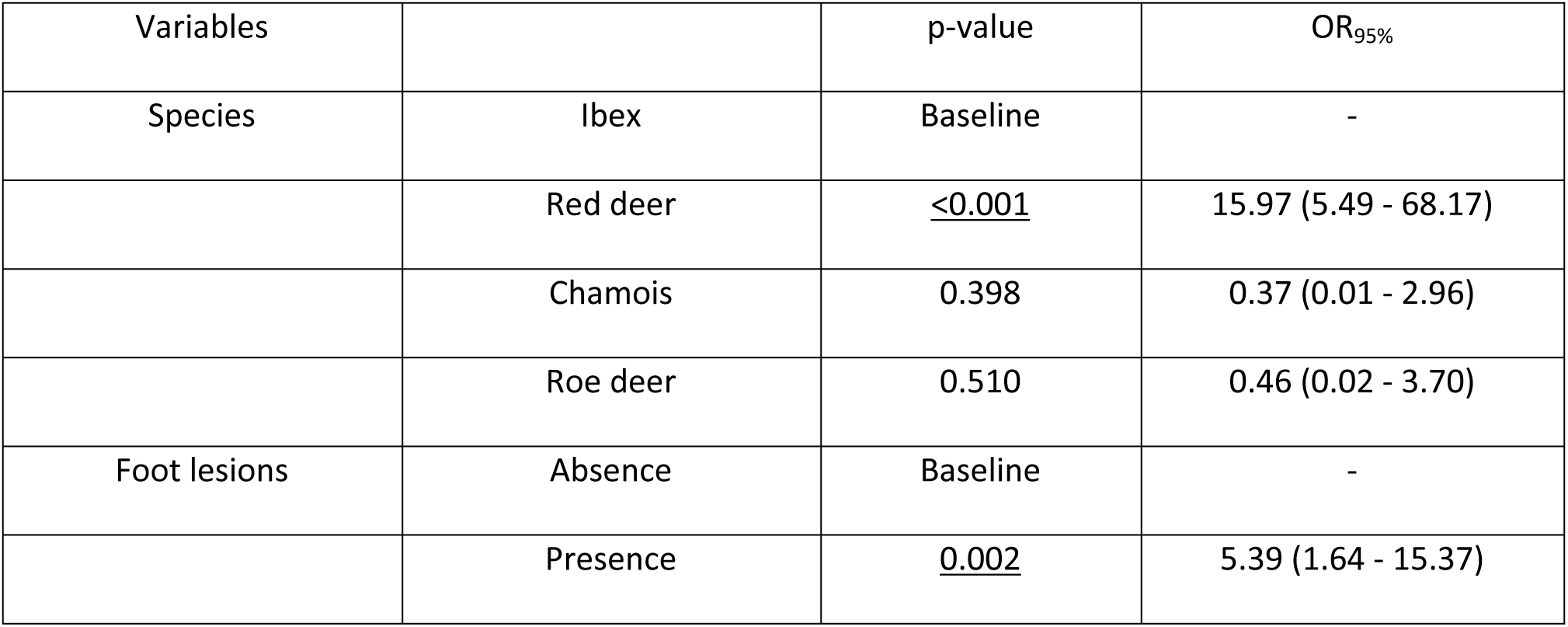
Multivariable analysis of risk factors for infection with benign *Dichelobacter nodosus* in wild ungulates.

In a further univariable analysis, the prevalence of infection in the four wild indigenous ruminants (all species pooled; this study) was compared to that found in sheep and cattle from the domestic ungulate study (Ardüser et. al. 2019). The model revealed that compared to all indigenous wild ruminants both “cattle” and “sheep” were respectively 221.38 (95% CI 2.97-7.00) and 4.53 (95% CI 154.52-326.43) times more likely to be infected with the benign strain of *D. nodosus*.

## Discussion

In this study, the prevalence of *D. nodosus* infections in wild ungulates in Switzerland was assessed for the first time, more specifically in all four indigenous wild ruminant species. These results provide a baseline necessary for the planning of a nationwide footrot control program in domestic livestock in Switzerland. Although the number of investigated animals was too low to provide strong data on a local level (e.g. cantonal), the sample size fulfilled the criteria for prevalence estimation per species on a country level. The occurrence of interspecies interactions among wild and domestic ruminants in an Alpine environment was additionally documented. Importantly, the study was conducted in parallel to a nationwide survey on infections with *D. nodosus* in possibly susceptible domestic livestock species The comparison of these data together with the reported interspecies interactions is of crucial importance in providing information on disease dynamics at the wildlife-livestock interface. (31).

### *Dichelobacter nodosus*-associated lesions

Two adult male Alpine ibex, the one infected with the benign and the other with the virulent strain, showed severe lesions consistent with footrot. These findings are in line with previous reports documenting that *D. nodosus* infections with both the benign and the virulent strain can be associated with severe foot lesions in Alpine ibex that are comparable to typical lesions of footrot in sheep (1,4). The classification into benign and virulent strains was developed in the framework of a study conducted in sheep and may only be applicable to sheep (9,32). The occurrence of severe lesions associated with both strains in ibex but not in other wild and domestic species suggest the existence of a species-specific difference in disease susceptibility, as it occurs for other pathogens such as *Mycoplasma conjunctivae* in wild (with disease signs) *vs* domestic Caprinae (without signs) and *Brucella abortus* in North American elk (*Cervus canadensis*) and Bison (*Bison bison*) (with signs) *vs* cattle (without signs) (4,33–35). As for the other two Alpine ibex that harbored the benign strain without foot lesions, it cannot be ruled out that they may have been in an early stage of the infection, before clinical disease signs developed. Nevertheless, it is not known whether infection with the benign strain requires the contribution of other factors (e.g. severe traumatic foot lesions, reduced fitness of the animal, co-infection with other microorganisms) for lesions to develop. Two red deer with reported foot lesions tested positive for the benign strain of *D. nodosus*. Whether these lesions were footrot-like could not be confirmed because neither the affected feet nor photographs were submitted for veterinary evaluation. However, *D. nodosus* has been already isolated from sole ulcers in farmed red deer in New Zealand (35). Although strain differentiation was not performed, it is likely that it was the benign type based on proteolytic effects observed in bacterial culture and because subsequent experimental infection of sheep resulted in the development of mild footrot lesions only. Infectious hoof diseases with similar clinical findings have been reported in cattle and free-ranging reindeer (36–38) involving pathogens such as *Fusobacterium necrophorum*, as well as *Treponema* sp. in cattle. Infection with *Treponema* sp. was also recently shown to be associated with a dramatic outbreak of severe foot lesions including osteomyelitis in free-ranging North American elk (*Cervus canadensis*) (39,40). Therefore, since all samples of the present study were tested via qPCR for the presence of *D. nodosus* only, we cannot exclude that other pathogens causing foot disease in domestic and wild ruminants may have been involved in the development of the observed lesions.

The presence of foot lesions was identified as a factor associated with the infection of benign *D. nodosus*. Obviously, the reported lesions were rather the consequence of the infection than a risk factor. This result from the risk factor analysis is also questionable because these lesions were only confirmed as footrot-like in the ibex.

### Interspecies interactions

Only 68% of the respondents answered (partially or fully) the questions regarding interspecies interactions on the data sheet. Additionally, considering that wildlife is most often active at dawn or dusk (6,41,42) it is therefore possible that the frequency and intensity of reported interactions was underestimated. Furthermore, because these data rely on reports of numerous voluntary participants, they may be biased in other ways. Mainly, the delivered information relies on memorized events of the participants and may have resulted in a lack of reporting or in misreporting. Nevertheless, this method represents the only feasible and efficient way of recruiting such data on a large geographical scale, and reported interspecies interactions at the wildlife/domestic animal interface were in agreement with previous findings (19,20). Most frequent interactions between wild and domestic ungulates involved cattle or sheep. Such data are useful to understand the epidemiology of infectious diseases in natural habitats, which are shared during certain periods by multiple species, wild and domestic.

### Identifying maintenance hosts

The present study revealed a very low prevalence of infection in all considered wild ungulate species. All prevalence calculated for wild species were significantly lower than those reported in both cattle (benign strain) and sheep (benign and virulent strain)(13). There is no given threshold of prevalence to conclude that a pathogen is maintained in a population or not, pathogen persistence in a population has been demonstrated to rely on the critical community size of a given population considering that the number of susceptible individuals does not drop under a certain threshold (43). Although in smaller wildlife populations it has been previously reported that some infections agents can persist with low incidences, provided that pathogens can circulate between subpopulations of several species and recurrent interspecific transmission occurs (33,44). However, in the case of *D. nodosus* the difference in prevalence between wild and domestic ungulates is striking. Additional points to consider are: 1) the endemic disease status of the Swiss sheep population (3,13); 2) the recently gained knowledge on the widespread occurrence of *D. nodosus* infection in asymptomatic livestock in Switzerland (88% of cattle positive with the benign strain)(3,13); 3) the much larger sizes of cattle and sheep populations compared to wild ungulate populations (24,45); 4) the more continuous distribution of domestic than wild animals over the country (ibex in particular occur in fairly isolated colonies(24,40); 5) the more frequent and intense intraspecific contacts among domestic animals and mixing of herds (grazing, shows, markets) and movements throughout the country (alpine summering, commercial exchanges); 6) the absence of healthy carriers for the virulent strain in wild ungulates; and 7) the reported interspecies interactions at the domestic animal/wildlife interface regarding mainly sheep and cattle. Therefore taking into consideration the previously mentioned points together with the comparison of the domestic ungulate study, our results suggest that sheep are maintenance hosts for both virulent and benign *D. nodosus* and cattle for the benign strain only. Thus, both likely represent the main source of infection for domestic and wild ungulates. Regarding cattle, the prevalence of infection with the benign strain is in agreement with reports from Norway and Australia (11,46). By contrast, our results suggest that wild ruminants are occasional spillover hosts without epidemiological significance. Nevertheless, the sample size used in this study does not allow to exclude potential differences in prevalence of infection on a local level. For example, the occurrence of previous outbreaks of footrot in wildlife, wildlife population management and densities, frequency and intensity of interspecific interactions on summer pastures and livestock practices such as already existing sanitation programs (virulent strain) are all factors that likely vary among regions.

Regarding the benign strain, infections were detected in at least one animal per wild ungulate species, with red deer showing the highest prevalence of infection (TP 6.08%). Risk factor analysis indicated that red deer are at a significantly higher risk of being infected with benign *D. nodosus* than ibex. This is in line with the fact that interspecies interactions involving red deer are most commonly observed with cattle (this study; 20) and that the prevalence of benign *D. nodosus* is highest in cattle (AP 83.3%; 13). It is therefore most likely that cattle act as source of infection for susceptible wild ungulates, as previously suspected (4). Since the upcoming nationwide control program will only focus on virulent *D. nodosus* in sheep, it is unclear whether it will influence the prevalence of benign *D. nodosus* in any species, i.e. the control program is not expected to significantly decrease the risk of footrot outbreaks caused by the benign strain of *D. nodosus* in ibex populations in the future.

## Conclusion

This study delivers crucial information for the design of the upcoming nationwide control program of virulent *D. nodosus* in sheep. Furthermore, the simultaneous, harmonized investigation of both wild and domestic species contributes to a better understanding of the epidemiology of footrot in Switzerland and will help targeting disease prevention measures, as it revealed that combatting the virulent strain does not necessitate to involve wildlife. The obtained data suggest that infections in wild ruminants are sporadic despite the high prevalence’s of infection in sheep and cattle and despite the widespread occurrence of interactions between wildlife and livestock. Overall, wildlife seems to be an incidental spillover host and not a maintenance host that may infect healthy sheep or re-infect sanitized herds. Since footrot lesions in ibex were also associated with the benign strain and the upcoming control program will only focus on the virulent strain in sheep, this program is not expected to influence disease occurrence in ibex in the future, especially when taking into consideration the high prevalence of benign *D. nodosus* in the Swiss cattle population. Further research is needed to evaluate the impact of footrot in ibex and whether it would be appropriate to take disease management actions to prevent outbreaks.

## Acknowledgements

We are grateful to all game wardens, hunters and cantonal hunting offices who contributed to the sample collection. We also thank all students and FIWI collaborators, especially Stefania Vannetti and Simone R. R. Pisano for their contributions to sample collection and processing. Many thanks go to the staff of the Institute of Bacteriology, especially Simon Feyer and Anita Jaussi, for their contributions to laboratory analyses. This study was supported by a grant of the Food Safety and Veterinary Office (FSVO) (1.17.f) with additional support from the Swiss Federal Office for Environment and the hunting authorities of the cantons of Fribourg, Grisons, Ticino, Schwyz and Nidwalden.

## Supporting information captions

**S1 Table. Cantons with PCR positive animals.** Estimated true**/apparent* prevalence of *D. nodosus*, with corresponding 95% confidence intervals indicated in parentheses and number of animals (positive/tested).

## References

1. Belloy L, Giacometti M, Boujon P, Waldvogel A. Detection of *Dichelobacter nodosus* in wild ungulates (*Capra ibex ibex* and *Ovis aries musimon*) and domestic sheep suffering from foot rot using a two-step polymerase chain reaction. Journal of Wildlife Diseases. 2007 Jan 1;43(1):82–8.

2. Rogdo T, Hektoen L, Slettemeås J, Jørgensen H, Østerås O, Fjeldaas T. Possible cross-infection of *Dichelobacter nodosus* between co-grazing sheep and cattle. Acta Veterinaria Scandinavica. 2012;54(1):19.

3. Zingg D, Steinbach S, Kuhlgatz C, Rediger M, Schüpbach-Regula G, Aepli M, et al. epidemiological and economic evaluation of alternative on-farm management scenarios for ovine footrot in Switzerland. Front Vet Sci. 2017; doi:10.3389/fvets.2017.00070.

4. Wimmershoff J, Ryser-Degiorgis M, Marreros N, Frey J, Romanens P, Gendron K, et al. Outbreak of severe foot rot associated with benign *Dichelobacter nodosus* in an Alpine ibex colony in the Swiss Prealps. Schweiz Arch Tierheilkd. 2015 May 5;157(5):277–84.

5. Volmer K, Hecht W, Weiß R, Grauheding D. Treatment of foot rot in free-ranging mouflon (*Ovis gmelini musimon*) populations—does it make sense? Eur J Wildl Res. 2008 Oct 1;54(4):657–65.

6. Meile P, Giacometti M, Ratti P. Der Steinbock: Biologie und Jagd. Salm; 2003. 269 p.

7. Kennan RM, Han X, Porter CJ, Rood JI. The pathogenesis of ovine footrot. Veterinary Microbiology. 2011 Nov;153(1–2):59–66.

8. Frosth S, König U, Nyman A-K, Pringle M, Aspán A. Characterisation of *Dichelobacter nodosus* and detection of *Fusobacterium necrophorum* and *Treponema* spp. in sheep with different clinical manifestations of footrot. Veterinary Microbiology. 2015 Aug;179(1–2):82–90.

9. Stäuble A, Steiner A, Normand L, Kuhnert P, Frey J. Molecular genetic analysis of *Dichelobacter nodosus* proteases AprV2/B2, AprV5/B5 and BprV/B in clinical material from European sheep flocks. Veterinary Microbiology. 2014 Jan;168(1):177–84.

10. Green LE, George TRN. Assessment of current knowledge of footrot in sheep with particular reference to *Dichelobacter nodosus* and implications for elimination or control strategies for sheep in Great Britain. The Veterinary Journal. 2008 Feb 1;175(2):173–80.

11. Knappe-Poindecker M, Gilhuus M, Jensen TK, Vatn S, Jørgensen HJ, Fjeldaas T. Cross-infection of virulent *Dichelobacter nodosus* between sheep and co-grazing cattle. Veterinary Microbiology. 2014 Jun;170(3–4):375–82.

12. Jubb, Kennedy & Palmer’s Pathology of Domestic Animals: Volume 1. St. Louis, Missouri: Elsevier; 2016. pp. 642–645.

13. Ardüser F, Moore-Jones G, Gobeli Brawand S, Dürr S, Steiner A, Ryser-Degiorgis M-P, et al. Dichelobacter nodosus in sheep, cattle, goats and South American camelids in Switzerland-Assessing prevalence in potential hosts in order to design targeted disease control measures. Prev Vet Med. 2019 May 6;104688.

14. Zuba JR. Hoof disorders in nondomestic artiodactylids. In: Fowler’s Zoo and Wild Animal Medicine. St. Louis, Missouri: Elsevier; 2012. pp. 619–27.

15. Delétraz C. Le piétin chez les ongulés sauvages: Etude clinique et épidémiologique chez le bouquetin des Alpes. DVM Thesis, Ecole Nationale Vétérinaire de Lyon, Lyon, France. 2002.

16. Egerton J., Roberts DS. Vaccination against ovine foot-rot. Journal of Comparative Pathology. 1971 Apr;81(2):179–85.

17. Locher I, Greber D, Holdener K, Lüchinger R, Haerdi-Landerer C, Schüpbach-Regula G, et al. Longitudinal *Dichelobacter nodosus* status in 9 sheep flocks free from clinical footrot. Small Ruminant Research. 2015 Nov;132:128–32.

18. Ryser-Degiorgis M-P, Bischof DF, Marreros N, Willisch C, Signer C, Filli F, et al. Detection of *Mycoplasma conjunctivae* in the eyes of healthy, free-ranging Alpine ibex: possible involvement of Alpine ibex as carriers for the main causing agent of infectious keratoconjunctivitis in wild Caprinae. Vet Microbiol. 2009 Mar 2;134(3–4):368–74.

19. Ryser-Degiorgis M-P, Ingold P, Tenhu H, Less AMT, Ryser A, Giacometti M. Encounters between Alpine ibex, Alpine chamois and domestic sheep in the Swiss Alps. Hystrix It J Mamm. 2002 Dec 20;13 (1–2).

20. Casaubon J, Vogt H-R, Stalder H, Hug C, Ryser-Degiorgis M-P. Bovine viral diarrhea virus in free-ranging wild ruminants in Switzerland: low prevalence of infection despite regular interactions with domestic livestock. BMC Vet Res. 2012 Oct 29;8:204.

21. 2013_nutztiere_schlussbericht.pdf [Internet]. [cited 2019 Feb 19]. Available from: http://www.alpfutur.ch/src/2013_nutztiere_schlussbericht.pdf.

22. Gortazar C, Diez-Delgado I, Barasona JA, Vicente J, De La Fuente J, Boadella M. The wild side of disease control at the wildlife-livestock-human interface: a review. Front Vet Sci. 2015 Jan; 14;1. doi:10.3389/fvets.2014.00027.

23. APHAEA project partners, Sonnenburg J, Ryser-Degiorgis M-P, Kuiken T, Ferroglio E, Ulrich RG, et al. Harmonizing methods for wildlife abundance estimation and pathogen detection in Europe—a questionnaire survey on three selected host-pathogen combinations. BMC Veterinary Research. 2016 Dec; 13(1). doi:10.1186/s12917-016-0935-x.

24. Jagdstatistik [Internet]. [cited 2019 Jun 20]. Available from: https://www.uzh.ch/wild/static/jagdstatistik/?page=wildtiere&dt=1&gr=1&tr=1&th=1&ca=CH&co=CH&caco=1&ys=2009&ye=2016&lang=de.

25. Greber D, Locher I, Kuhnert P, Butty M-A, Holdener K, Frey J, et al. Pooling of interdigital swab samples for PCR detection of virulent *Dichelobacter nodosus*. Journal of Veterinary Diagnostic Investigation. 2018 Mar;30(2):205–10.

26. Stewart DJ, Claxton PD. Ovine footrot. Clinical diagnosis and bacteriology. Australian Standard Diagnostic Techniques for Animal Diseases. LA Corner and TJ Bagust. Melbourne, Standing Committee on Agriculture and Resource Management Sub-committee on Animal Health Laboratory Standards. CSIRO; 1993.

27. Reimoser F, Duscher T, Duscher A, Jenny H, Nigsh N. Rothirsch im Rätikon–drei Länder, drei Jagdsysteme, eine Wildart. Hohenems: Vorarlberger Jägerschaft, Amt für Umwelt Fürstentum Liechtenstein, Amt für Jagd und Fischerei Graubünden. 2015;

28. Andersen R, Duncan P, Linnell JDC. The European roe deer: the biology of success. Scandinavian University Press; 1998. 384 p.

29. Baumann M, Muggli J, Thiel D, Thiel-Egenter C, Thürig M, Volery P, et al. Jagen in der Schweiz; Auf dem Weg zur Jagdprüfung. Salm Verlag, Wohlen; 2012.

30. Rogan WJ, Gladen B. Estimating prevalence from the results of a screening test. Am J Epidemiol. 1978 Jan;107(1):71–6.

31. Martin C, Pastoret P-P, Brochier B, Humblet M-F, Saegerman C. A survey of the transmission of infectious diseases/infections between wild and domestic ungulates in Europe. Veterinary Research. 2011;42(1):70.

32. Niggeler A, Tetens J, Stäuble A, Steiner A, Drögemüller C. A genome-wide significant association on chromosome 2 for footrot resistance/susceptibility in Swiss white Alpine sheep. Animal Genetics. 2017;48(6):712–5.

33. Gelormini et al. Infectious keratoconjunctivitis in wild Caprinae: merging field observations and molecular analyses sheds light on factors shaping outbreak dynamics. BMC Veterinary Research. 2017; doi:10.1186/s12917-017-0972-0.

34. Godfroid J. Brucella spp. at the wildlife-livestock interface: an evolutionary trajectory through a livestock-to-wildlife “host jump”? Vet Sci. 2018 Sep 18; 5(3). doi:10.3390/vetsci5030081.

35. Olsen SC, Johnson C. Comparison of abortion and infection after experimental challenge of pregnant bison and cattle with *Brucella abortus* strain 2308. Clin Vaccine Immunol. 2011 Dec 1;18(12):2075–8.

36. Klitgaard K, Boye M, Capion N, Jensen TK. Evidence of multiple Treponema phylotypes involved in bovine digital dermatitis as shown by 16S rRNA gene analysis and fluorescence in situ hybridization. J Clin Microbiol. 2008 Sep;46(9):3012–20.

37. Nielsen MW, Strube ML, Isbrand A, Al-Medrasi WDHM, Boye M, Jensen TK, et al. Potential bacterial core species associated with digital dermatitis in cattle herds identified by molecular profiling of interdigital skin samples. Veterinary Microbiology. 2016 Apr;186:139–49.

38. Handeland K, Boye M, Bergsjø B, Bondal H, Isaksen K, Agerholm JS. Digital necrobacillosis in Norwegian wild tundra reindeer (*Rangifer tarandus tarandus*). Journal of Comparative Pathology. 2010 Jul 1;143(1):29–38.

39. Clegg SR, Mansfield KG, Newbrook K, Sullivan LE, Blowey RW, Carter SD, et al. Isolation of digital dermatitis treponemes from hoof lesions in wild North American elk (*Cervus elaphus*) in Washington State, USA. Fenwick BW, editor. Journal of Clinical Microbiology. 2015 Jan;53(1):88–94.

40. Han S, Mansfield KG. Severe hoof disease in free-ranging Roosevelt elk (*Cervus elaphus roosevelti*) in southwestern Washington, USA. Journal of Wildlife Diseases. 2014 Apr;50(2):259–70.

41. Göran Cederlund. Activity patterns in moose and roe deer in a north boreal forest. Holarctic Ecology. 1989;12(1):39–45.

42. Georgii B, Schröder W. Home range and activity patterns of male red deer (*Cervus elaphus L.*) in the alps. Oecologia. 1983 May;58(2):238–48.

43. Bartlett MS. Measles periodicity and community size. Journal of the Royal Statistical Society Series A (General). 1957;120(1):48–70.

44. Almberg ES, Cross PC, Smith DW. Persistence of canine distemper virus in the Greater Yellowstone Ecosystem’s carnivore community. Ecological Applications. 2010;20(7):2058–74.

45. Schweizer Bauernverband - der Dachverband der Schweizer Landwirtschaft - Schweizer Bauernverband [Internet]. [cited 2019 Jun 21]. Available from: https://www.sbv-usp.ch/de/

46. Laing EA, Egerton JR. The occurrence, prevalence and transmission of *Bacteroides nodosus* infection in cattle. Res Vet Sci. 1978 May;24(3):300–4.

